# The model arbuscular mycorrhizal fungus *Rhizophagus irregularis* harbours endosymbiotic bacteria with a highly reduce genome

**DOI:** 10.1101/2021.09.13.460061

**Authors:** Romain Savary, Frédéric G. Masclaux, Ian R. Sanders

**Affiliations:** Laboratory for Biological Geochemistry, School of Architecture, Civil and Environmental Engineering, Ecole Polytechnique Fédérale de Lausanne (EPFL), 1015 Lausanne, Switzerland; Department of Ecology and Evolution, Biophore Building, University of Lausanne, 1015 Lausanne, Switzerland; Vital-IT Group, SIB Swiss Institute of Bioinformatics, 1015 Lausanne, Switzerland

## Abstract

Arbuscular mycorrhizal fungi (AMF; Glomeromycotina) are symbionts of most plant species that are known to possess unique intracytoplasmic endosymbiotic bacteria with an enigmatic role. *Candidatus* Moeniiplasma glomeromycotorum (*Ca*Mg) was shown to be widespread along the AMF phylogeny and present in most AMF species and isolates of those species. The model AMF species, *Rhizophagus irregularis*, that can be cultivated *in vitro* and for which a lot of genomic information now exists, would be the ideal model to study the true nature of the *Ca*Mg-AMF symbiosis. However, *R. irregularis* was never found to host endobacteria. Here we show by DNA sequencing that *R. irregularis* can, indeed, host *Ca*Mg (*Ri-Ca*Mg). However, this appears rare as only one *R. irregularis* isolate out of 58 hosted *Ca*Mg. In that isolate, the endosymbiotic bacterial population was genetically homogenous. By sequencing the complete genome of the bacteria, we found that its genome is among the smallest of all known *Ca*Mg and Mycoplasma-like genomes, with a highly reduced gene repertoire, suggesting a strong adaptation to the intracellular life. We discuss our findings in the light of previous literature on *Ca*Mg and on the same AMF isolates and suggest that these endosymbionts are more likely parasites than non-obligatory mutualists.

## Results and Discussion

Arbuscular mycorrhizal fungi (AMF) are symbionts of most plant species, exchanging essential soil nutrients for photosynthates. This symbiosis is central to terrestrial ecosystems, as AMF diversity influences the diversity of plant communities, as well as their structure and productivity (van der Heijden *et al*., 1998). Arbuscular mycorrhizal fungi (AMF, Glomeromycotina) are known to possess unique intracytoplasmic endosymbiotic bacteria, that exhibit the typical reduced-gene repertoire of host-dependent symbionts (Naito *et al*., 2015). Two main species of AMF endobacteria have been described. The first, *Candidatus* Glomeribacter gigasporum (*Ca*Gg) has only been described in the *Gigasporaceae* family and was classified as a member of the *Burkholderia* (Bianciotto *et al*., 2003, Mondo *et al*., 2012). The second, *Candidatus* Moeniiplasma glomeromycotorum (*Ca*Mg, or previously called *Mycoplasma*-related endobacteria, MRE, Naito *et al*., 2017), was found to be present in nearly the entire Glomeromycotina phylogeny, including *Gigasporaceae* (Naumann *et al*., 2010, Desirò *et al*., 2014, Toomer *et al*., 2015) and are related to a number of animal and plant pathogenic bacteria within the *Mycoplasma* genus (Naumann *et al*., 2010). Only few AMF species, such as *Rhizophagus irregularis* were found to be free of these bacteria (Naumann *et al*., 2010, Toomer *et al*., 2015, Torres-Cortés *et al*., 2015). To date, the nature of the relationship between *Ca*Mg and its AMF host is still unclear. Kuo (2015) suggested that in order to understand the functions of *Ca*Mg in the symbiosis with AMF, it would be critical to study the AMF hosts and their genomes. Comparing genomes of AMF species that are able, and those that are unable, to form the symbiosis with endosymbiont bacteria could give insights into this relationship. Lacking the ability to form such associations *R. irregularis* was, thus, suggested as offering a rare opportunity to understand the symbiosis (Kuo, 2015). This AMF species, in comparison with other AMF species that associate with *Ca*Mg, could potentially reveal what type of functions were lost or gained following the loss of this endosymbiont. Indeed, larger genomics and transcriptomics resources are available for this model AMF species compared to any other AMF species (Tisserant *et al*., 2013, Ropars *et al*., 2016, Wyss *et al*., 2016, Savary *et al*., 2018).

An even better experimental design to understand the true nature of these bacteria would be to be able to compare between individuals of the same species, with or without endobacteria. Despite a large literature on *R. irregularis* and *Ca*Mg (Naumann *et al*., 2010), the presence of *Ca*Mg within an *R. irregularis* isolate has never been reported. Identifying *R. irregularis* isolates that are able to form and maintain this symbiosis, along with some others that are not able, would be a first step in paving the way to understanding the true nature of the interaction. Indeed, it would provide the ideal comparative model as *R. irregularis* together with the large genomics resources available is probably the easiest AMF species to propagate and maintain in *in-vitro* cultures with the largest number of isolates in collections (Lin *et al*., 2014).

In this study, we provide evidence that *R. irregularis* is able to maintain a symbiosis with *Ca*Mg. We provide the first sequence *Rhizophagus irregularis-Ca*Mg (*Ri-Ca*Mg) genome assembly and a draft sequence of the associated *Rhizophagus irregularis* genome. We, thus, provide the ideal model to understand what allows this symbiosis to occur and whether it is beneficial or detrimental to the host. Finally, we discuss the possible parasitic nature of *Ca*Mg and its wider ecological role in the light of previous literature.

### *Ca*Mg occurs in *R. irregularis* populations

To confirm the presence or absence of endobacteria in *R. irregularis* isolates from highly diverse geographical origins, we analysed previously published ddRAD-seq data on 58 isolates of *R. irregularis*, 20 isolates of some closely-related species, namely *Rhizophagus intraradices*, *Rhizophagus proliferus* and two isolates (LPA8, CH3) of the genetically closest species to *R. irregularis; Rhizophagus* sp. LPA8-CH3 (Wyss *et al*., 2016, Savary *et al*., 2018, Table S1). We screened for sequences mapping to either the *Ca*Mg genome or *Ca*Gg genome. The ddRAD-seq method is a non-selective method of sequencing, allowing the sequencing of any potential holobionts associated with the AMF. As all ddRAD-seq data originated from *in-vitro* AMF cultures (Savary *et al*., 2018), detection of bacterial reads belonging specifically to *Ca*Mg or *Ca*Gg is highly probable if the bacteria are present in one of the isolates and should be easy to map. Also due to the sterile nature of the cultures, there is also a low probability of sequences from other contaminant bacteria. An advantage of the available data was that there were multiple replicates of each isolate. Each replicate represents material collected from one Petri dish of a culture. While the detection of a high number of reads mapping to *Ca*Mg genomes in one replicate might be considered as a contamination, a consistent high mapping of reads in all replicates, to one or several *Ca*Mg genomes, would strongly suggest the presence of a *Ca*Mg symbiont.

We were able to detect *Ca*Mg sequences in *Rhizophagus irregularis* (*Ri-Ca*Mg), as well as in the closely related species to *R. irregularis, Rhizophagus* sp. LPA8-CH3 (*R.-Ca*Mg, Fig. 1a, Table S1, see Savary et al., 2018). However, only one *R. irregularis* isolate (A2), out of 58 isolates, contained a significant number of reads mapping to the *Ca*Mg genome. Interestingly, no reads mapped the isolates of the two closely-related species of *R. irregularis*, namely *R. intraradices* and *R. proliferus* suggesting that *Ca*Mg are rare in population of *Rhizophagus* species. Reads of the other isolate (CH3) belonging to the enigmatic closely-related species to *R. irregularis*, i.e., *R*. sp. LPA8-CH3 only mapped to the *Rhizophagus clarus-Ca*Mg (*Rc-Ca*Mg) genome, but no other *Ca*Mg genomes confirmed the presence of an endobacteria in CH3 isolate (Fig. 1a). As expected, sequences mapping to the *Gigasporaceae* specific endobacteria (*Ca*Gg) were not found in any of the *Rhizophagus* species.

**Figure 1.**
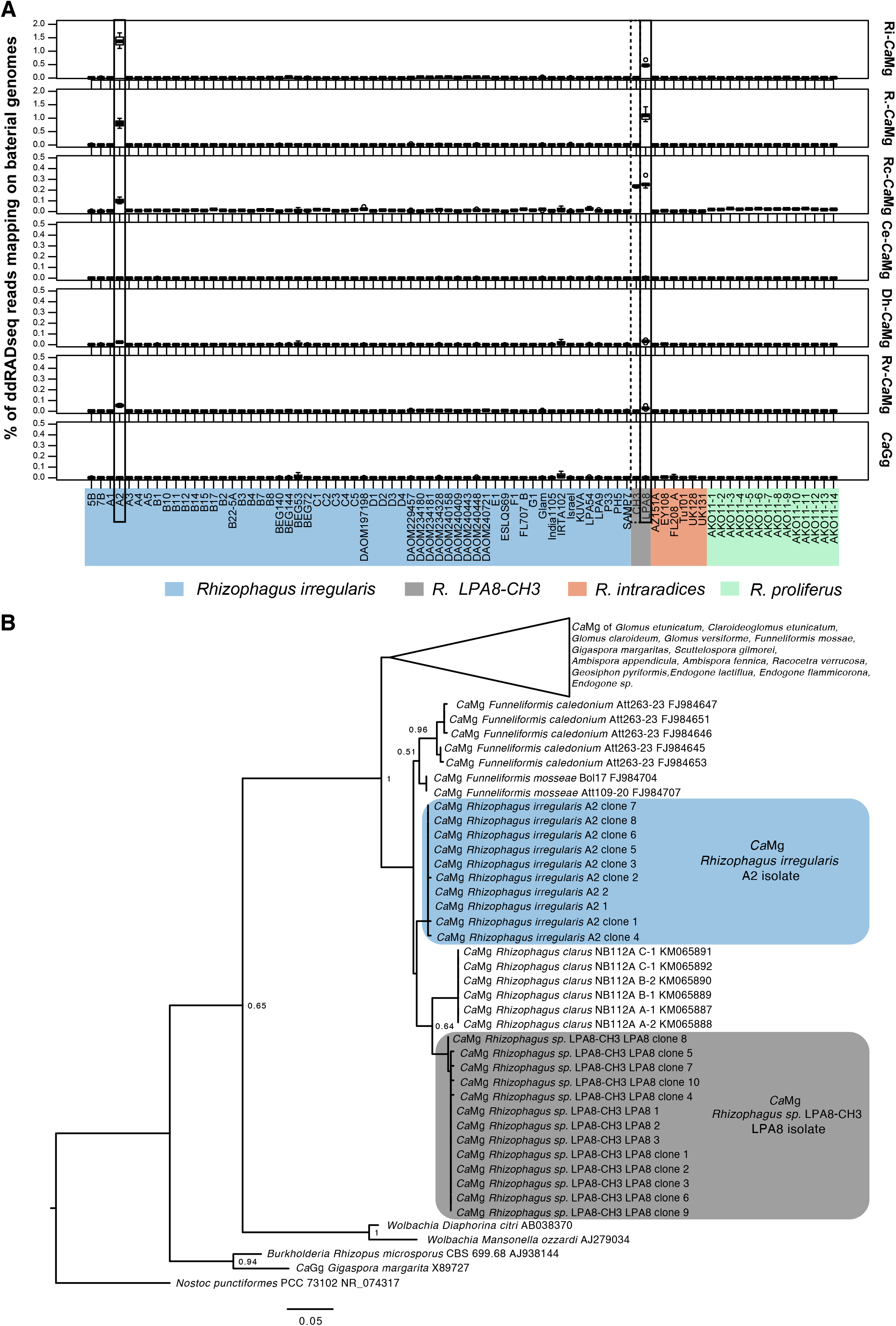
**a)** Boxplot of the percentage of ddRAD-seq reads from each of the 58 *R. irregularis* isolates, 20 other isolates from known *Rhizophagus* species and 2 isolates of the closest species to *R. irregularis* (3-5 ddRAD-seq replicates per isolates) that mapped to four published *Ca*Mg genomes (*Rhizophagus clarus-Ca*Mg, *Claroideoglomus etunicatum-Ca*Mg, *Dentiscutata heterogama-Ca*Mg, *Racocetra verrucose-Ca*Mg), as well as to new assemblies of *Ri-Ca*Mg and *R.-Ca*Mg generated in this study, and to the other known AMF endosymbiotic bacterial species *Ca*Gg. Percentages are calculated as the proportion of reads mapped to bacterial genomes compared to the total number of reads in a given sample. The colour corresponds to the different species (blue = *R. irregularis*, grey = *R. sp*. LPA8-CH3, red = *R. intraradices* and green = *R. proliferus*). **b)** Maximum likelihood phylogeny built with MEGA7 (Kumar *et al*., 2016) of *Ca*Mg 16S rRNA sequences and cloned sequences from this study and those obtained from publications. The NCBI accession number from previously published sequences can be found on the tree and new *Ca*Mg 16S rRNA sequences of this study can be found on NCBI with accession number MW161341-MW161363.

### *R. irregularis* harbours a homogenous population of *Ca*Mg

In order to: i) confirm the presence of the endosymbiont, ii) construct a phylogeny and iii) detect the within-isolate endosymbiotic bacterium genetic variation, all isolates were subjected to 16S rRNA amplification (see Supporting Information and Table S1). Isolates that showed *Ca*Mg specific amplification were cloned and sequenced. The 16S rRNA sequences allowed us to confirm the presence of *Ca*Mg in isolate A2 and LPA8, as expecting according to ddRAD-seq read mapping. It is known that some AMF species host several divergent haplotypes of *Ca*Mg within the cytoplasm, such as the 5 *Ce-Ca*Mg haplotypes, coexisting in *C. etunicatum* cytoplasm (Naito *et al*., 2015). In contrast, *R. clarus* and *R. verrucosa* both contain a single homogenous population of endobacterial haplotypes within the fungal cytoplasm (Naito *et al*., 2015). In the case of *Ri-Ca*Mg and *R.-Ca*Mg, the population was also shown to be homogenous (Fig. 1b) based on 8 and 10 cloned 16S rRNA sequences of *Ri-Ca*Mg and *R.-Ca*Mg, respectively. This suggests that the *Ca*Mg population was not enriched by horizontal haplotype transfer in these fungi as proposed for several other AMF species (Toomer *et al*., 2015).

A genetically homogenous endobacterial population, present in only some individuals is consistent with non-obligate mutualism (Toomer *et al*., 2015). Moreover, the *Ri-Ca*Mg and *R.-Ca*Mg 16S rRNA sequences revealed a high relatedness to *Ca*Mg found in the other closely-related AMF species*, Rhizophagus clarus, Funneliformis caledonium* and *Funneliformis mossae* (Fig. 1b). This suggests that in these species, *Ca*Mg were vertically inherited. Indeed, similar trends were observed using a gene fragment encoding a metallo-beta-lactamase family protein (MBLFP, Toomer et al., 2015, Supporting Information Fig. S1). Similarly, the *Rc-Ca*Mg genome (bacteria present in *Rhizophagus clarus*, i.e., the closest AMF to *Rhizophagus irregularis* from which a *Ca*Mg was sequenced) was the *Ca*Mg to which the highest percentage of *Ca*Mg reads from isolates A2 and LPA8 mapped (Naito *et al*., 2015, Fig. 1a). The presence of a potential *Ca*Mg in the isolate CH3 (species; *R. sp* LPA8-CH3) could not be confirmed since DNA amplification was not successful with either 16S rRNA or MBLFP primer pairs, despite several attempts.

### The smallest and largest *Ca*Mg genomes observed in AMF and the smallest recorded mycoplasma-like genome

To obtain the *Ri-Ca*Mg and *R*.-CaMg genome sequences, we jointly extracted AMF and *Ca*Mg DNA from the respective *in-vitro* AMF cultures and sequenced their meta-genomes. Three A2 and three LPA8 independant *in-vitro* culture plates were extracted to obtain four replicates of the *Ri-Ca*Mg genome (A2_3, A2_4, A2_6, A2_WG) and three replicates of *R.-Ca*Mg genome (LPA8_1, LPA8_3, LPA8_4, Table S2 and Table S3 and Supporting Information Fig. S4). Assemblies are available as Newly assembled genome of CaMg.zip and Newly assembled genome hosting CaMg.zip. Surprisingly, meta-genome sequencing, assembly, analysis and comparison with other available *Ca*Mg and Mycoplasma genomes (see Supporting Information) revealed that the two genomes of *Rhizophagus* endobacteria spanned the two extremes in size of known *Ca*Mg endobacterial and Mycoplasma genomes (Fig. 2). Despite that phylogenetic analysis of 16S rRNA revealed high relatedness between the two *Ca*Mg found in the two closely related *Rhizophagus* species, there has been major expansion/loss and rearrangement between these two genomes (Fig. 2, Supporting Information, Fig. S2), confirming the large genome plasticity found in these species (Naito *et al*., 2015, Kuo, 2015). The percentage GC content was maintained across all *Ca*Mg as the two newly reported *Ca*Mg genomes fell within the natural *Ca*Mg GC% range (32.1% in *Rc-Ca*Mg to 34.3% in *Ce-Ca*Mg), with approximately 32.5% in *R.-Ca*Mg and 33% in *Ri-Ca*Mg. *Ri-Ca*Mg was well assembled, with a high similarity between the four replicate assemblies (Supporting Information Fig. S3). The replicate with the largest genome size (A2_3) contained only 7 contigs and a total size of 565 759bp (Supporting Information Fig. S3). This makes *Ri-Ca*Mg the smallest *Ca*Mg genome sequenced to date and, thus, the smallest known AMF endosymbiont. It is also the smallest reported mycoplasma-like genome, as *Mycoplasma genitalium* is slightly larger (580’070 bp, Fraser *et al*., 1995). The gene number is highly reduced with only 492 CDS (Prokka annotation)/ 571 CDS (GeneMarks annotation) and highly similar to the *M. genitalium* gene content (489/536). In the *R.-Ca*Mg genome, the number of genes is three times higher (1513/1661) with similar gene density. *Ri-Ca*Mg also showed the smallest gene density with 863-900 genes per Mb. Using BUSCO (Simão *et al*., 2015) and the Tenericute database, we detected low Complete BUSCO percentages for the present, and all previously published *Ca*Mg (1.8% for *Ri-Ca*Mg and 2.4% for *DhCa*Mg) genomes, while all Mycoplasma genomes had a high score (>90%), (Table S2). Similar results were found using the BUSCO Bacteria database (Table S2). The reduction in gene number, the low BUSCO score and the large rearrangement (Supporting Information Fig. S2) highlight the extreme plasticity in these highly related genomes and the loss of a large number of genes as a result of an extreme adaptation to an endosymbiotic way of life.

**Figure 2.**
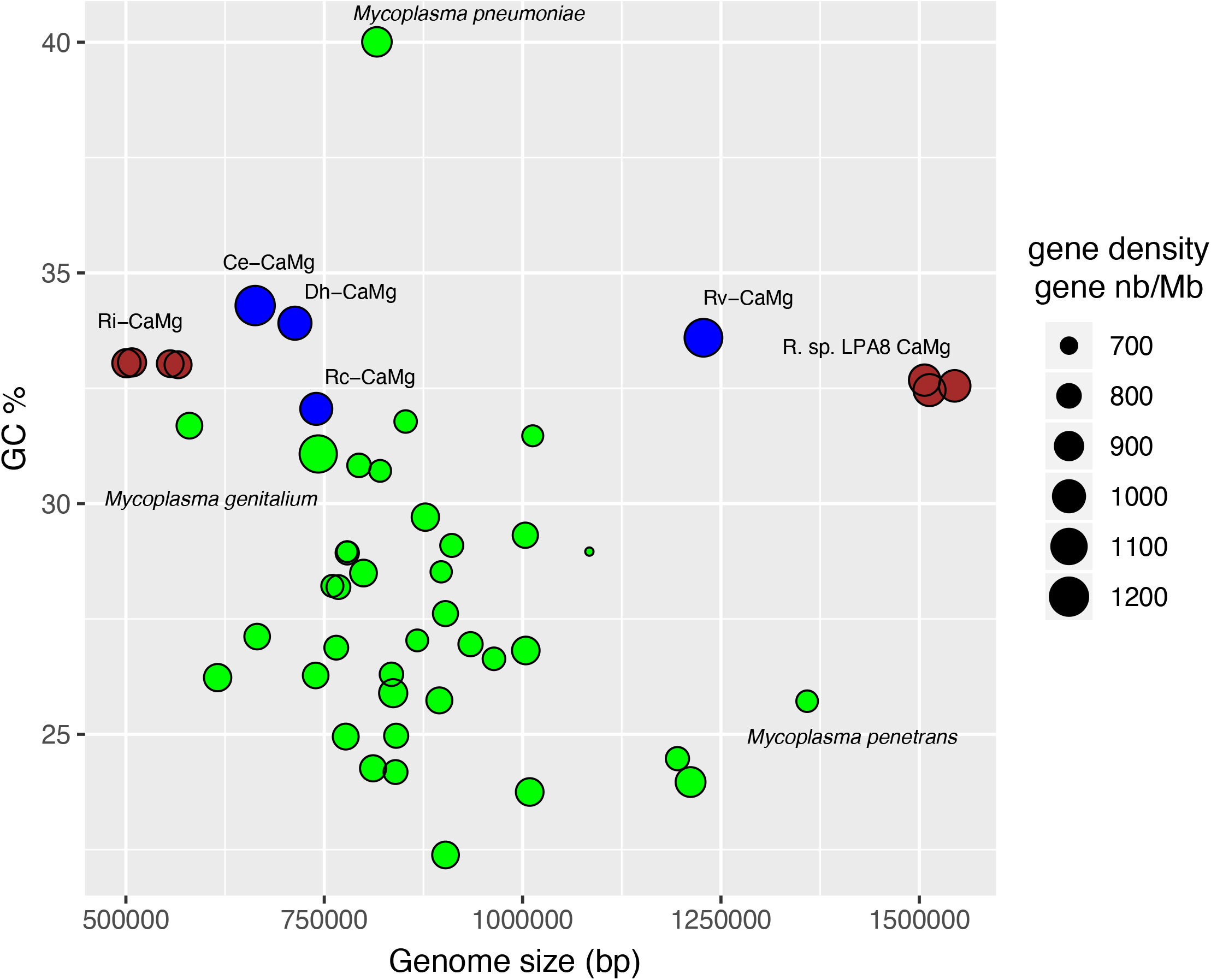
Genome size and GC% of 38 Mycoplasma (green) and 6 *Ca*Mg genomes. Previously sequenced *Ca*Mg in blue, and the two *Ca*Mg genomes sequenced in this study in brown with 4 replicates of *Ri-Ca*Mg (*R. irregularis*, A2 isolate) and 3 replicates of *R.-Ca*Mg (*R. sp*. LPA8-CH3, LPA8 isolate, Savary *et al*., 2018). Circle size corresponds to the gene density i.e. number of genes per million base pairs. The number of genes found in the genome come from the Prokka annotation.

### The potential role of *RiCa*Mg bacteria in *R. irregularis*

It has been hypothesized that endobacteria played a major role in the evolution of AMF, as they have been maintained internally and transmitted for more than 100 million years across different AMF lineages (Torres-Cortés *et al*., 2015). It was, therefore, suggested that they probably confer an increase of fitness to their host (Naumann *et al*., 2010). This was confirmed for the endosymbiotic bacteria *Ca*Gg, in *G. margarita*, whose presence resulted in higher sporulation rates, bioenergetic capacity and ATP production (Salvioli *et al*., 2016). In contrast, Toomer *et al*., 2015 suggested that *Ca*Mg are parasites, rather than mutualists, as they harbour a high intra-host genetic diversity which is typical of parasites that maintain genetic diversity through recombination and horizontal transmission. In contrast, we showed that *Ca*Mg living in *R. irregularis* do not harbor high within host diversity compare to other *Ca*Mg based on their 16S sequences and have reduced genome size and gene content. Results from previous studies on *R. irregularis* isolate A2 have shown a strong decrease in sporulation rate (Koch *et al*., 2004) and low colonization rate (Savary *et al*., 2020) in this *R. irregularis* isolate compared to other non-*Ca*Mg harbouring isolates of the same species and population. These results suggest that the presence of the endobacteria within this isolate could potentially come at a fitness cost to the A2 fungus and supports the hypothesis of a parasitic nature of the *Ca*Mg (Toomer *et al*., 2015). Curing isolate A2 of its endosymbiont will be a key step in order to better understand the true nature of the interactions between AMF and *Ca*Mg (Salvioli *et al*., 2016). A further indication of the possible parasitic nature of these bacteria in *R. irregularis* is that they were only found in one *R. irregularis* isolate out of 58 isolates implying that other *R. irregularis* have lost the symbiont by unknown means. Indeed, *in-vitro* culturing might have been one of the means leading to this rarity of *Ca*Mg, however the presence of the endobacteria in the isolate A2, that was subcultured in *in-vitro* for more than 20 years suggests that this does not affected their presence. In mutualistic symbioses, the increased fitness afforded by hosting a mutualistic symbiont is expected to lead to all individuals possessing the symbiont. This was clearly not the case in *R. irregularis*. The ability of this particular AMF clade to get rid of these potential endoparasites could have been one evolutionary innovation which contributes to their ecological success compared to other AMF clades that still harbour endobacteria. Indeed, *R. irregularis* and close relatives are cosmopolitan and often present in large quantities in a variety of different ecosystems around the world (Davison *et al*., 2015).

### A role in of the tripartite symbiosis?

A high variation in AMF phenotypic traits and plant responses have been widely observed, even when using AMF isolates of the same species (Koch *et al*., 2006, Klironomos *et al*., 2003). This variation could be the result of different AMF genetic background (Savary *et al*., 2018, Savary *et al*., 2020). However, the role of the presence/absence of a potential endoparasitic bacteria in AMF used for plant-AMF experiment has never been studied and could be an interesting and potential source of variation in the outcome of the symbiosis. The system we are presenting could be suitable to test the role of AMF endobacteria in the tripartite symbiosis and its wider role in ecology as these *Ca*Mg bacteria are widely spread in the AMF genetic tree.

## Supporting information

Newly assembled genome hosting CaMG

Newly assembled genome of CaMG

## Acknowledgement

We would like to thank Chanz Robbins for the helpful comments on the manuscript and Jeremy Bonvin for the help in fungal culturing. This study was funded by the Swiss National Science Foundation (Grant number: 31003A_162549 to IRS). Sanger sequences can be found on NCBI under accession number MW161341-MW161363 for 16S rRNA sequences. Raw meta-sequencing data of the two AMF isolates, A2 and LPA8 and their respective *Ca*Mg, with all replicates and their assembled genomes can be found under BioProject number PRJNA670049.

## Author contributions

RS planned and designed the research. RS performed experiments and analysed the data with the help of FGM. RS and IRS wrote the manuscript.

## Supporting information

**Figure S1**

Maximum-likelihood phylogeny of the *Ca*Mg MBLFP sequences

**Figure S2**

Multiple genome alignment of the best assembled replicates of *Ri-Ca*Mg (replicate 3) and *R.-Ca*Mg LPA8 (replicate 3)

**Figure S3**

Multiple genome alignment of the A2 *Ri-Ca*Mg replicates

**Figure S4**

Blobplot of assembled contigs for all metagenomes in each replicate

**Table S1**

List of AMF isolates (origin, species) tested in this study for presence or absence of CaMg

**Table S2**

Summary of *Ca*Mg genome assemblies

**Table S3**

Summary of AMF genome assemblies hosting *Ca*Mg

## Notes

### Competing Interest Statement

The authors have declared no competing interest.

## References

Bianciotto V, Lumini E, Bonfante P, Vandamme P. 2003. ‘Candidatus Glomeribacter gigasporarum’ gen. nov., sp nov., an endosymbiont of arbuscular mycorrhizal fungi. International Journal of Systematic and Evolutionary Microbiology 53: 121–124.

Davison J, Moora M, Ouml, pik M, Adholeya A, Ainsaar L, Ba A, Burla S, Diedhiou AG, Hiiesalu I, et al. 2015. Global assessment of arbuscular mycorrhizal fungus diversity reveals very low endemism. Science 349(6251): 970–973.

Desiro A, Salvioli A, Ngonkeu EL, Mondo SJ, Epis S, Faccio A, Kaech A, Pawlowska TE, Bonfante P. 2014. Detection of a novel intracellular microbiome hosted in arbuscular mycorrhizal fungi. Isme Journal 8(2): 257–270.

Fraser CM, Gocayne JD, White O, Adams MD, Clayton RA, Fleischmann RD, Bult CJ, Kerlavage AR, Sutton G, Kelley JM, et al. 1995. The minimal gene complement of mycoplasma-genitalium. Science 270(5235): 397–403.

Klironomos JN. 2003. Variation in plant response to native and exotic arbuscular mycorrhizal fungi. Ecology 84(9): 2292–2301.

Koch AM, Croll D, Sanders IR. 2006. Genetic variability in a population of arbuscular mycorrhizal fungi causes variation in plant growth. Ecology Letters 9(2): 103–110.

Koch AM, Kuhn G, Fontanillas P, Fumagalli L, Goudet J, Sanders IR. 2004. High genetic variability and low local diversity in a population of arbuscular mycorrhizal fungi. Proceedings of the National Academy of Sciences of the United States of America 101(8): 2369–2374.

Kombrink A, Tayyrov A, Essig A, Stockli M, Micheller S, Hintze J, van Heuvel Y, Durig N, Lin C-w, Kallio PT, et al. 2019. Induction of antibacterial proteins and peptides in the coprophilous mushroom Coprinopsis cinerea in response to bacteria. Isme Journal 13(3): 588–602.

Kuo C-H. 2015. Scrambled and not-so-tiny genomes of fungal endosymbionts. Proceedings of the National Academy of Sciences of the United States of America 112(25): 7622–7623.

Lin K, Limpens E, Zhang ZH, Ivanov S, Saunders DGO, Mu DS, Pang EL, Cao HF, Cha HH, Lin T, et al. 2014. Single Nucleus Genome Sequencing Reveals High Similarity among Nuclei of an Endomycorrhizal Fungus. Plos Genetics 10(1).

Mondo SJ, Toomer KH, Morton JB, Lekberg Y, Pawlowska TE. 2012. Evolutionary stability in a 400-million-year-old heritable facultative mutualism. Evolution 66(8): 2564–2576.

Naito M, Desiro A, Gonzalez JB, Tao G, Morton JB, Bonfante P, Pawlowska TE. 2017. ‘Candidatus Moeniiplasma glomeromycotorum’, an endobacterium of arbuscular mycorrhizal fungi. International Journal of Systematic and Evolutionary Microbiology 67(5): 1177–1184.

Naito M, Morton JB, Pawlowska TE. 2015. Minimal genomes of mycoplasma-related endobacteria are plastic and contain host-derived genes for sustained life within Glomeromycota. Proceedings of the National Academy of Sciences of the United States of America 112(25): 7791–7796.

Naumann M, Schussler A, Bonfante P. 2010. The obligate endobacteria of arbuscular mycorrhizal fungi are ancient heritable components related to the Mollicutes. Isme Journal 4(7): 862–871.

Ropars J, Toro KS, Noel J, Pelin A, Charron P, Farinelli L, Marton T, Kruger M, Fuchs J, Brachmann A, et al. 2016. Evidence for the sexual origin of heterokaryosis in arbuscular mycorrhizal fungi. Nature Microbiology 1(6).

Salvioli A, Ghignone S, Novero M, Navazio L, Venice F, Bagnaresi P, Bonfante P. 2016. Symbiosis with an endobacterium increases the fitness of a mycorrhizal fungus, raising its bioenergetic potential. Isme Journal 10(1): 130–144.

Savary R, Dupuis C, Masclaux FG, Mateus ID, Rojas EC, Sanders IR. 2020. Genetic variation and evolutionary history of a mycorrhizal fungus regulate the currency of exchange in symbiosis with the food security crop cassava. Isme Journal 14(6): 1333–1344.

Savary R, Masclaux FG, Wyss T, Droh G, Corella JC, Machado AP, Morton JB, Sanders IR. 2018. A population genomics approach shows widespread geographical distribution of cryptic genomic forms of the symbiotic fungus Rhizophagus irregularis. Isme Journal 12(1): 17–30.

Simao FA, Waterhouse RM, Ioannidis P, Kriventseva EV, Zdobnov EM. 2015. BUSCO: assessing genome assembly and annotation completeness with single-copy orthologs. Bioinformatics 31(19): 3210–3212.

Tisserant E, Malbreil M, Kuo A, Kohler A, Symeonidi A, Balestrini R, Charron P, Duensing N, Frey NFD, Gianinazzi-Pearson V, et al. 2013. Genome of an arbuscular mycorrhizal fungus provides insight into the oldest plant symbiosis. Proceedings of the National Academy of Sciences of the United States of America 110(50): 20117–20122.

Toomer KH, Chen XH, Naito M, Mondo SJ, den Bakker HC, VanKuren NW, Lekberg Y, Morton JB, Pawlowska TE. 2015. Molecular evolution patterns reveal life history features of mycoplasma-related endobacteria associated with arbuscular mycorrhizal fungi. Molecular Ecology 24(13): 3485–3500.

Torres-Cortes G, Ghignone S, Bonfante P, Schussler A. 2015. Mosaic genome of endobacteria in arbuscular mycorrhizal fungi: Transkingdom gene transfer in an ancient mycoplasma-fungus association. Proceedings of the National Academy of Sciences of the United States of America 112(25): 7785–7790.

van der Heijden MGA, Klironomos JN, Ursic M, Moutoglis P, Streitwolf-Engel R, Boller T, Wiemken A, Sanders IR. 1998. Mycorrhizal fungal diversity determines plant biodiversity, ecosystem variability and productivity. Nature 396(6706): 69–72.

Wyss T, Masclaux FG, Rosikiewicz P, Pagni M, Sanders IR. 2016. Population genomics reveals that within-fungus polymorphism is common and maintained in populations of the mycorrhizal fungus Rhizophagus irregularis. Isme Journal 10(10): 2514–2526.

