## Supporting Information for "The model arbuscular mycorrhizal fungus *Rhizophagus irregularis* harbours endosymbiotic bacteria with a highly reduce genome"

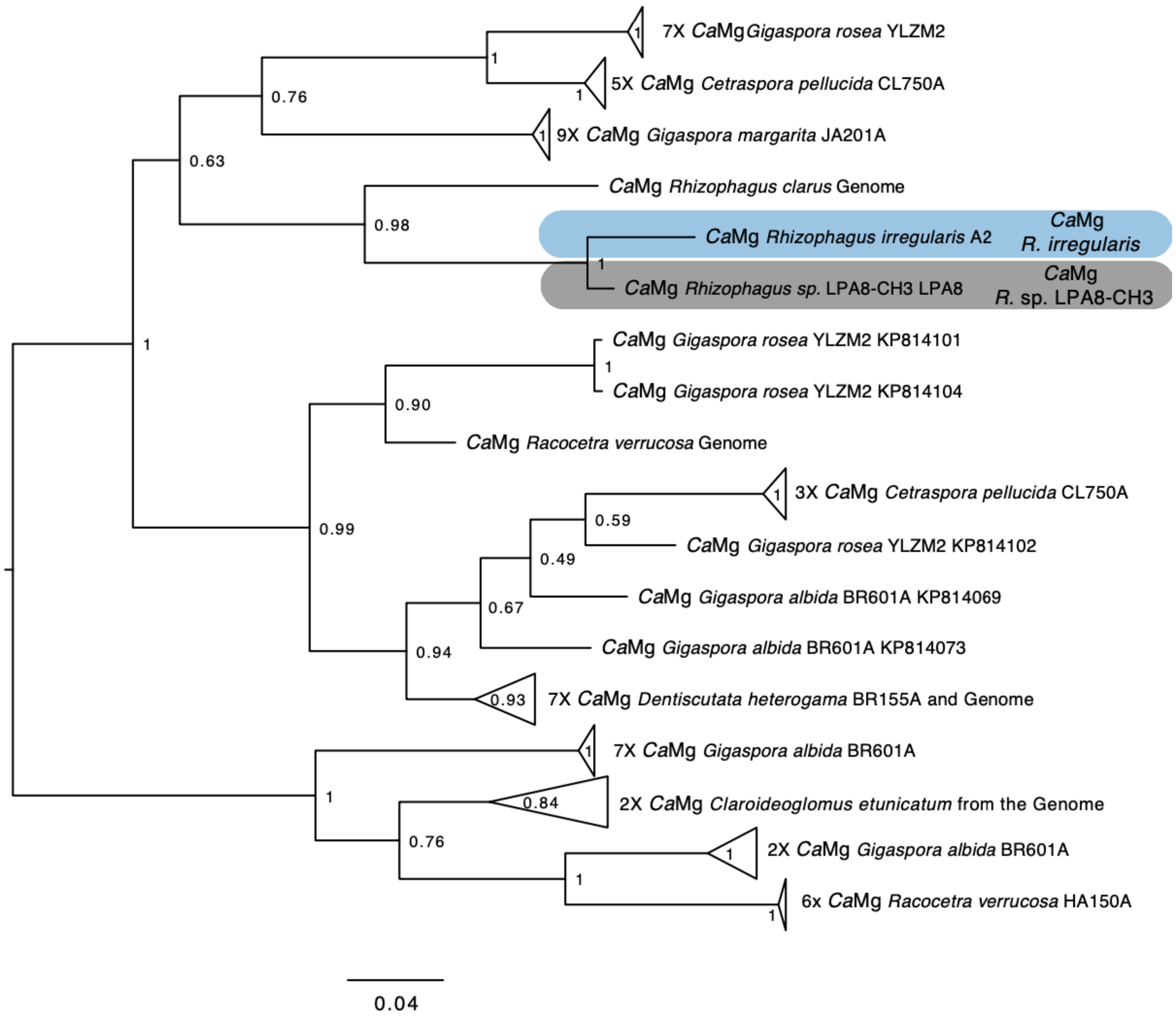

**Fig. S1** Maximum-likelihood phylogeny mid-point rooted build with MEGA7 (Kumar et al., 2016) of the *Ca*Mg gene fragment encoding a metallo-beta-lactamase family protein (MBLFP). MBLFP sequences obtained from Toomer *et al.,* 2015 are included as well as the MBLFP sequences of four sequenced *Ca*Mg genomes. NCBI accession numbers can be found next to each sequence.

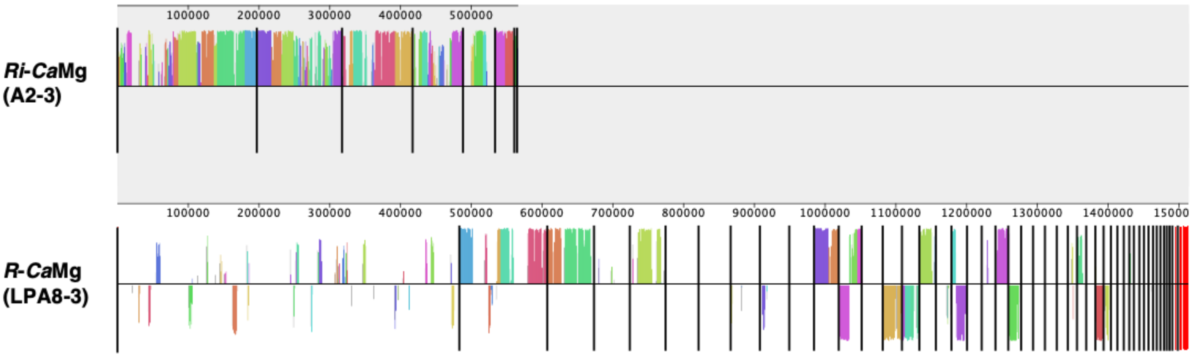


**Fig. S2** Multiple genome alignment of the best assembled replicates of *Ri-Ca*Mg (replicate 3) and *R.-Ca*Mg LPA8 (replicate 3) performed with MAUVE (Darling et al., 2004). Colour coding represents conserved regions between the genomes. Black bar shows the limits of the scaffolds. Scaffolds are ordered by size.

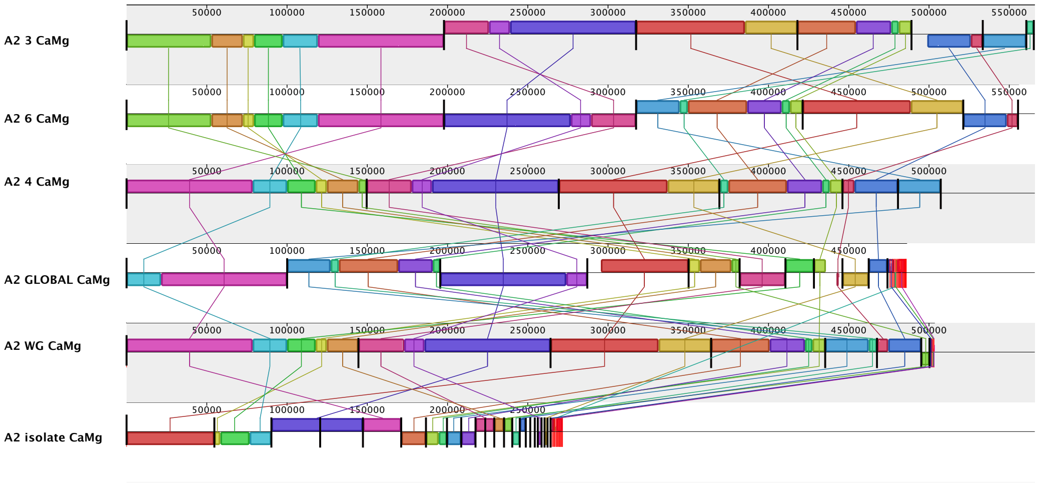


**Fig. S3** Multiple genome alignment of the A2 *Ri-Ca*Mg replicates performed with MAUVE (Darling et al., 2004), where the numbered suffix following the isolate name represents the replicate number. The additional sample name suffix GLOBAL or ALL corresponds to the assembly performed with the reads of all replicates pooled. The sample with the suffix "isolate" corresponds to the assembly of an *RiCa*Mg genome obtained from bacterial DNA isolated from the R. irregularis-A2 DNA. Colours represent similar regions between genomes. Black bars show the limits of the scaffolds. Scaffolds are ordered by size.

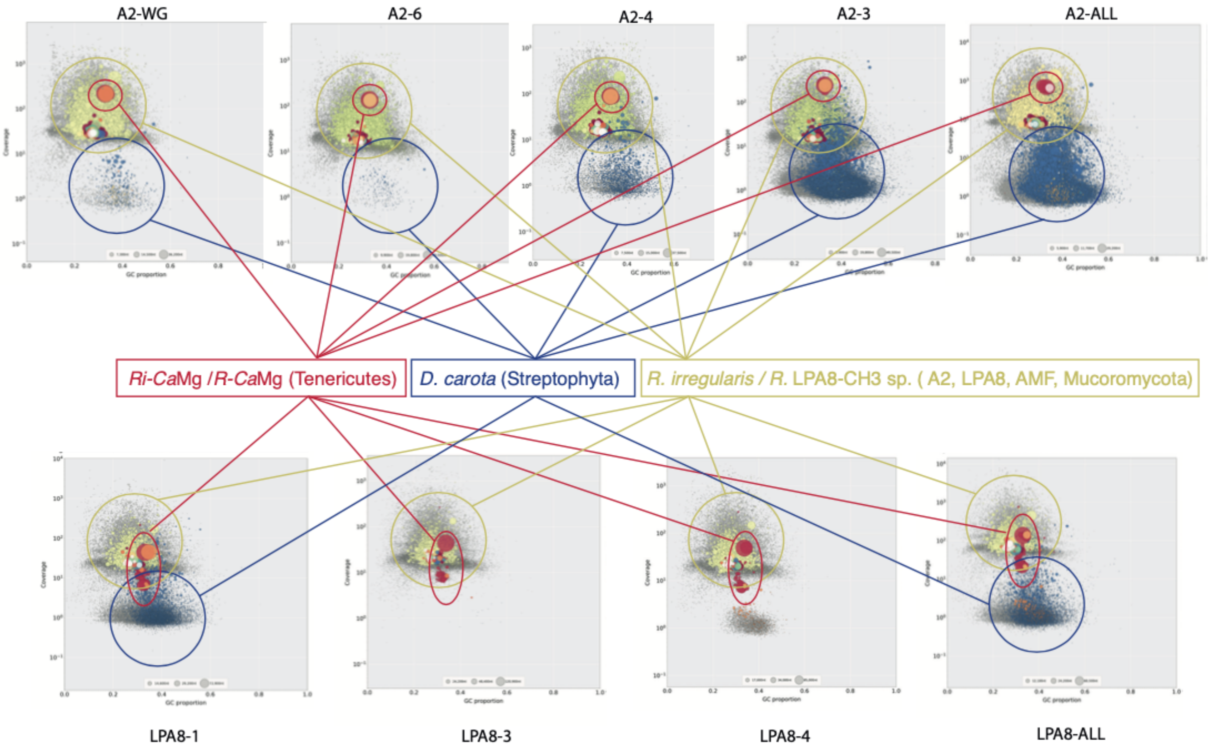


**Fig. S4** Blobplot of assembled contigs for all metagenomes in each replicate, where the numbered suffix represents the replicate number and the suffix ALL corresponds to the assembly performed with the reads of all replicates pooled.

**Supporting Methods:**

*ddRAD-seq and workflow*

Published double digest restriction-site associated DNA (**ddRADseq**) data on a population of 80 *R. irregularis* isolates, and related species, from the bioprojects PRJNA268659 and PRJNA326895 (Wyss *et al.,* 2016, Savary *et al.,* 2018), were used to detect the presence of *Ca*Mg reads within the data. Read quality check and trimming was carried out following Savary et *al.,* 2018. The demultiplexed reads of each isolate, and in each of the three to five replicates, were mapped using the Novoalign software (Novocraft-Technologies, 2014) to the *DhCa*Mg genome, an *Ca*Mg present in *Dentiscutata heterogama* (Torres-Cortés *et al.,* 2015) and on *CeCa*Mg (*Claroideoglomus etunicatum Ca*Mg), *RcCa*Mg (*Rhizophagus clarus Ca*Mg) and *RvCa*Mg (*Racocetra verrucosa Ca*Mg) genomes (Naito *et al.,* 2015) as well as to two newly sequenced genomes, *Ri-Ca*Mg and *R. sp.* LPA8-CH8 *Ca*Mg, described below. Reads were also mapped to endobacteria found in *Gigaspora margarita*, *Ca*Gg (*Candidatus* Glomeribacter gigasporum). SAM files were converted to BAM files and indexed with Samtools (Li *et al.,* 2009). Visual inspection of mapped reads was performed using IGV (Robinson *et al.,* 2011). Summary tables generated by Novoalign with the number of mapped reads per replicate were used to calculate a percentage of bacterial reads per samples presented in the boxplot in Fig. 1a.

*DNA extraction and 16S rRNA and MBLFP gene amplification*

Sterile, two-compartment plate cultures were produced for all 80 *R. irregularis* isolates and related species and grown for at least 3 months to obtain sufficient spores for DNA extraction. DNA extraction was performed as described in Savary et *al.,* 2018. The 16S rRNA gene portion was amplified using primers 109F (109F-1 and 109F-2) and 1184R (1184R-1, 1184R-2 and 1184R-3, Naumann *et al.,* 2010). The MBLFP gene was amplified using MBLFP18f and MBLFP644r primers (Toomer *et al.,* 2015). A new reverse primer for the 16S rRNA (16SrRNA-RcCaMg: 5'-AGTTACCTTGGCAGTCTGC-3') was designed on the basis of the *Rc*-CaMg 16S rRNA sequence specific to its genome (Naito *et al.,* 2015). Similarly, a new forward (MBLFP-RcCaMg-f, 5'-GAAAYYGGAGAAAAACTGAYYTAGYYAA-3') and reverse (MBLFP-RcCaMg-r, 5'-GARGCATGMARTAWKTCYTCT-3') degenerate primer set for MBLFP was designed, based on the homologue MBLFP gene found in the *Rc-Ca*Mg genome. The following protocol; a PCR mix of 23µl with Qiagen reagents (1X Buffer, 1.5mM $MgCl_2$, 0.2 mM dNTPs, 0.5 µM forward and reverse primer, 0.4 U of *Taq* polymerase) was applied to 2µl of 1 to 25ng/µl of DNA of each of the isolates. The amplification on a Biometra T1 Thermocycler PCR machine was performed under the following conditions: 5 min at 95°C followed by 35 cycles at 95°C for 10s, 50°C for 30s and 72°C for 2min, a final elongation step at 72°C for 10 min. A slight modification for MBLFP was necessary, by changing the annealing temperature to 49°C and by adding $MgCl_2$ for a final concentration of 2.25 mM. Purification and Sanger sequencing was performed by GATC biotech (Germany). Alignment and sequence cleaning was performed using MEGA 7 (M Kumar *et al.,* 2016).

*Localisation of* Ca*Mg*

To confirm the strict endocytoplasmic localisation of *Ca*Mg, DNA from AMF isolates suspected to be carrying *Ca*Mg based on ddRAD-seq mapping and Sanger sequencing, were re-extracted from sterile two- compartment plates under a UVC sterilized laminar hood. After dissolution of the fungal compartment for 30 min in citrate buffer (0.0062 M of citric acid anhydrous and 0.0028 M of sodium citrate tribasic dihydrate), the pellet of spores and hyphae were washed with $ddH_20$, which was used as PCR control. The pellet was surface sterilized with 2% chloramide-T for 3 min, and subsequently washed with 0.03% of Penicillin-streptomycin before being immersed in a 2mL Eppendorf tube filled with $ddH_20$. Each tube was sonicated for 1 min. The $ddH_20$ used for sonication served as a second control to detect potential contaminants or external *Ca*Mg during PCR. Clean spores were finally extracted with the DNeasy plant mini kit (QIAGEN). As a control, DNA from AMF isolates potentially having *Ca*Mg in their cytoplasm were extracted independently in sterile conditions. Water used for washing and sonication was included as controls during DNA extraction and PCR reactions. No PCR product was detected in this way from the washed water and the sonicated water in any of the PCRs, thus suggesting that no contaminant or *Ca*Mg are present outside the AMF host.

*Cloning and sequencing of the haplotypic population of* Ca*Mg*

*Ca*Mg were suggested to harbour a large diversity of haplotypes within one AMF isolate (Naito et *al.,* 2015). Thus, cloning of the 16S rRNA amplicon was performed with the StrataClone PCR Cloning Kit (Agilent Technologies®) for each isolate carrying an *Ca*Mg and for which we were able to amplify the 16S region.

*Phylogeny of 16S rRNA and MBLFP genes*

A maximum-likelihood phylogeny was inferred in MEGA7 with a bootstrapping method of 500 replicates (Kumar et *al.,* 2016) using 16S rRNA sequences obtained from Sanger sequencing as well as publicly available sequences (Naumann *et al.,* 2010, Toomer *et al.,* 2015, Naito *et al.,* 2015). A second phylogeny, performed using the same guidelines, was built with MBLFP sequences generated in this study along with previously published sequences (Toomer *et al.,* 2015) and homologue sequences present within the four different *Ca*Mg genomes. All trees were edited with Figtree v1.4.2 (Rambaut & Drummond, 2009) and Adobe Illustrator (2014.1.0 Release).

*Ca*Mg *genome sequencing*

We sequenced the genome of *Ca*Mg present in isolates A2 (*Rhizophagus irregularis*) and LPA8 (*Rhizophagus* sp.). We used AMF DNA obtained from the extraction of 3 months old split plates cultures. The whole genome sequencing of both the AMF host and its bacterial symbiont were performed by constructing libraries with the TruSeq Nano protocol (Illumina). We produced 4 libraries (A2-3, A2-4, A2-6, A2-WG) from 3 independent DNA extractions of A2

(3 dual-compartment culture plates, plate 3, plate 4, plate 6). The DNA extraction replicate 6 was used to build 2 libraries (A2-6, A2-WG). We produced 3 libraries from 3 independent DNA extractions (3 split plates) of LPA8. These libraries were paired-end sequenced on an Illumina sequencer Hiseq2500 for 100 nt.

*Genome assembly and annotation*

Sequences were cleaned of adapters and from low quality reads. All sequencing replicates were assembled independently with the metaSPAdes assembler (Nurk et al., 2017). Blobtools (Laetsch and Blaxter, 2017) was used to identify contigs blasting *Ca*Mg. Mean GC content and mean coverage of these contigs was also calculated from Blobtools outputs. Blobplots per host-symbiont combination (A2-*Ca*Mg and LPA8-*Ca*Mg) showing GC%, coverage for all contigs with their clade identity is found in Fig. S4. Only contigs identified as Tenericutes and contigs defined as Bacteria undefined and found in similar GC% and coverage as the Tenericutes contigs, where retained. Contigs smaller than 1000bp were removed. BUSCO was used to assess the quality of the assembly. Prokka (Seemann, 2014) and GeneMarkS (Besemer *et al.,* 2001) were used to annotate all genomes. A similar strategy using Blobtools was used to identify AMF contigs and to generate the draft genomes of A2 and LPA8. In these genomes, contigs smaller than 1000bp were as well removed.

### References


**Besemer J, Lomsadze A, Borodovsky M. 2001.** GeneMarkS: a self-training method for prediction of gene starts in microbial genomes. Implications for finding sequence motifs in regulatory regions. *Nucleic Acids Research* **29**(12): 2607-2618.

**Darling ACE, Mau B, Blattner FR, Perna NT. 2004.** Mauve: Multiple alignment of conserved genomic sequence with rearrangements. *Genome Research* **14**(7): 1394-1403.

**Kumar S, Stecher G, Tamura K. 2016.** MEGA7: Molecular Evolutionary Genetics Analysis Version 7.0 for Bigger Datasets. *Molecular Biology and Evolution* **33**(7): 1870-1874.

**Laetsch DR and Blaxter ML. 2017.** BlobTools: Interrogation of genome assemblies [version 1; peer review: 2 approved with reservations]. *F1000Research* 2017, **6**:1287

**Li H, Handsaker B, Wysoker A, Fennell T, Ruan J, Homer N, Marth G, Abecasis G, Durbin R, Genome Project Data P. 2009.** The Sequence Alignment/Map format and SAMtools. *Bioinformatics* **25**(16): 2078-2079.

**Naito M, Morton JB, Pawlowska TE. 2015.** Minimal genomes of mycoplasma-related endobacteria are plastic and contain host-derived genes for sustained life within Glomeromycota. *Proceedings of the National Academy of Sciences of the United States of America* **112**(25): 7791-7796.

**Naumann M, Schussler A, Bonfante P. 2010.** The obligate endobacteria of arbuscular mycorrhizal fungi are ancient heritable components related to the Mollicutes. *Isme Journal* **4**(7): 862-871.

**Novocraft-Technologies**. 2014. Novoalign. Available from http://www.novocraft.com.

**Nurk S, Meleshko D, Korobeynikov A, Pevzner PA. 2017.** metaSPAdes: a new versatile metagenomic assembler. *Genome Research* **27**(5): 824-834.

**Robinson JT, Thorvaldsdottir H, Winckler W, Guttman M, Lander ES, Getz G, Mesirov JP. 2011.** Integrative genomics viewer. *Nature Biotechnology* **29**(1): 24-26.

**Savary R, Masclaux FG, Wyss T, Droh G, Corella JC, Machado AP, Morton JB, Sanders IR. 2018.** A population genomics approach shows widespread geographical distribution of cryptic genomic forms of the symbiotic fungus Rhizophagus irregularis. *Isme Journal* **12**(1): 17-30.

**Seemann T. 2014.** Prokka: rapid prokaryotic genome annotation. *Bioinformatics* **30**(14): 2068-2069.

**Toomer KH, Chen XH, Naito M, Mondo SJ, den Bakker HC, VanKuren NW, Lekberg Y, Morton JB, Pawlowska TE. 2015.** Molecular evolution patterns reveal life history features of mycoplasma-related endobacteria associated with arbuscular mycorrhizal fungi. *Molecular Ecology* **24**(13): 3485-3500.

**Torres-Cortes G, Ghignone S, Bonfante P, Schussler A. 2015.** Mosaic genome of endobacteria in arbuscular mycorrhizal fungi: Transkingdom gene transfer in an ancient mycoplasma-fungus association. *Proceedings of the National Academy of Sciences of the United States of America* **112**(25): 7785-7790.

**Wyss T, Masclaux FG, Rosikiewicz P, Pagni M, Sanders IR. 2016.** Population genomics reveals that within-fungus polymorphism is common and maintained in populations of the mycorrhizal fungus Rhizophagus irregularis. *Isme Journal* **10**(10): 2514-2526.
